# Laser-activated drug implant for controlled release to the posterior segment of the eye

**DOI:** 10.1101/2020.06.17.111641

**Authors:** Xingyu He, Zheng Yuan, Samantha Gaeke, Winston W.-Y. Kao, S. Kevin Li, Daniel Miller, Basil Williams, Yoonjee C. Park

## Abstract

The current standard of care for posterior segment eye diseases, such as age-related macular degeneration and diabetic macular edema, is frequent intravitreal injections or sustained-release drug implants. Drug implants have side effects due to the burst release of the drugs, and their release cannot be easily controlled after implantation. Present study attempts to develop a dosage-controllable drug delivery implant which consists of a nanoporous biodegradable PLGA capsule and light-activated liposomes. Controllable drug release from the implant was achieved by using pulsed near-infrared (NIR) laser both in vitro and in vivo. The in vitro drug release kinetics from two different initial dose implants, 1000 μg and 500 μg, was analyzed by fitting zero order and first order kinetics, as well as the Korsmeyer-Peppas and Higuchi models. The 1000 μg and 500 μg implants fit the first-order and zero-order kinetics model, respectively, the best. The multiple drug releases in the vitreous was determined by in vivo fluorimeter, which was consistent with the in vitro data. The dose released was also clinically relevant. Histology and optical and ultrasound imaging data showed no abnormality in the eyes received implant treatment suggesting that the drug delivery system was safe to the retina. This on-demand dose-controllable drug delivery system could be potentially used for long-term posterior eye disease treatment.

## 1. Introduction

Posterior segment eye diseases such as age-related macular degeneration (AMD), diabetic macular edema (DME) and proliferative vitreoretinopathy (PVR) are serious chronic diseases that may result in vision loss. More than 100 million people around the world are suffering from these chronic posterior eye diseases.^1–2^ The current standard of care for posterior segment eye disease involves frequent repetitive intravitreal injections or sustained-release drug implants. Typically, intravitreal injections are administered monthly or every other month depending on disease and response to treatment. This is due to the relatively short pharmacokinetics of these medications, and there is a relatively short half-life of drugs delivered to the vitreous cavity (within hours – days).^3^ However, there is a significant burden to the patient, the patient’s family, and the health system because current intravitreal therapies require every 4 to 12 week administration over many years. To resolve the issue, intravitreally injectable sustained-release implants have been developed to prolong therapeutic efficacy.^4,5^ Biodegradable implants, such as Ozurdex®, used PLGA (poly lactide-co-glycolic acid) as a drug depot and designed for 6 month-sustained release. The drug release mechanism of Ozurdex implant relies entirely on degradation rate of a PLGA matrix where drug is impregnated. However, most of the drug is released initially after treatment within 1 month due to polymer degradation,^6^ leading to high initial drug concentrations in the vitreous or retina. This high initial dose or burst release of drug from the implant are considered to cause local side effects such as elevated intraocular pressure and cataract formation, etc.^7^

We previously developed a size-exclusive nanoporous biodegradable PLGA capsule as a dosage-controllable drug delivery implant, which was stable and safe in rabbit eyes for 6 months.^8^ In the present study, we combined the capsule with light-activatable drug-encapsulated liposomes, to create a light-activated dose-controllable implant. Our hypothesis is that light-activatable drug-encapsulated liposomes inside the nanoporous PLGA capsule release drug when irradiated by near-infrared laser irradiation and the drug diffuses through the nanoporous structure to the surrounding media (Figure 1). We focused on drug release from the implant by laser irradiation both in vitro and in vivo to demonstrate feasibility of using the implant as a dose-controllable drug delivery system to the posterior segment of the eye. Drug release kinetics in vitro were analyzed utilizing the implants with two different dosages.

**Figure 1.**
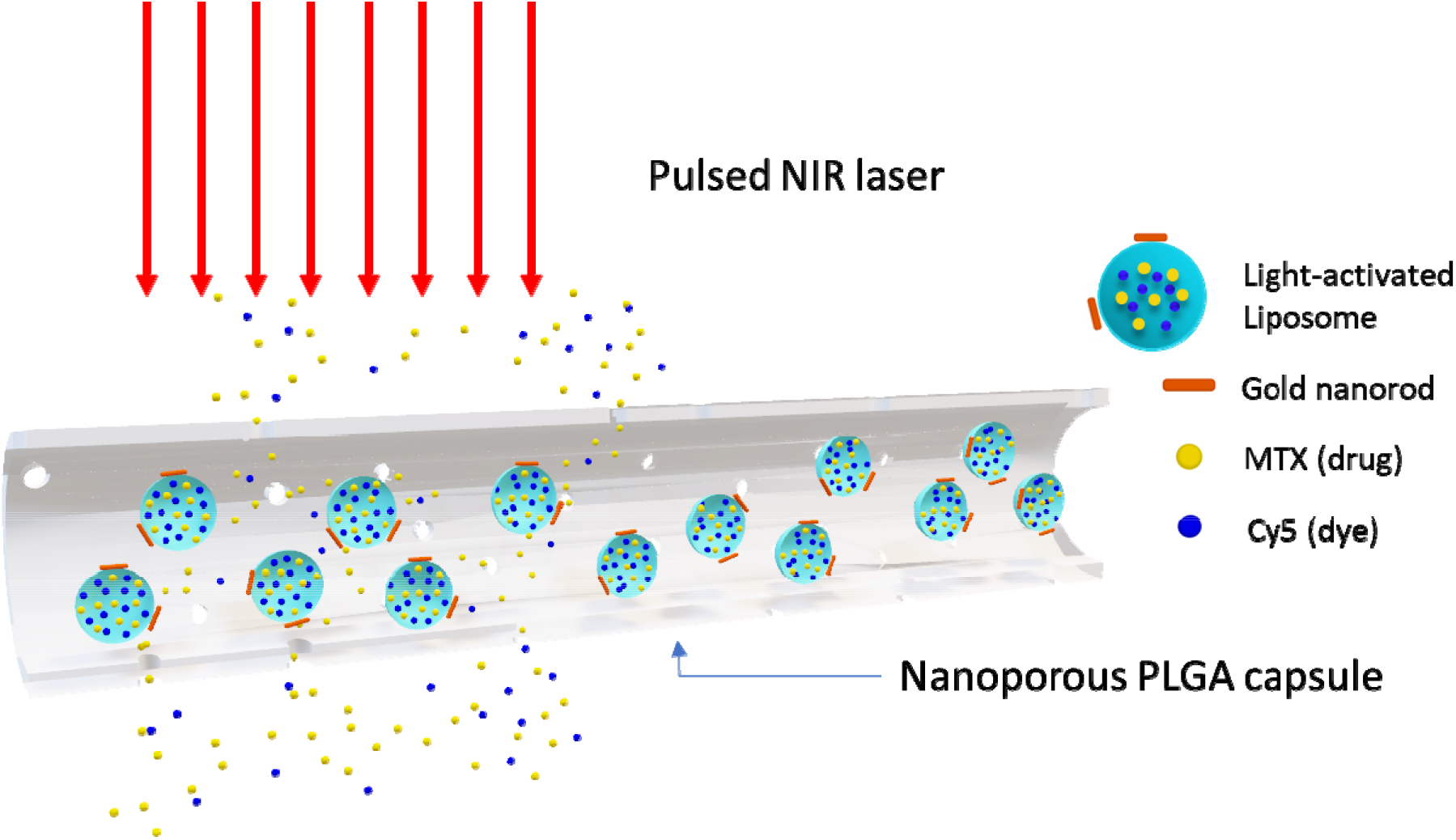
Schematic of light-activated drug implant under pulsed NIR laser irradiation. Drug and dye molecules are released from nanopores of the implant

The light-activatable liposomes co-encapsulate methotrexate (MTX) and Cy5 fluorescent dye. MTX is an anti-inflammatory drug, used to treat proliferative vitreoretinopathy (PVR) and non-infectious uveitis.^9^ Cy5 is used to quantitively analyze hydrophilic molecules released by laser trigger in vivo using fluorescence imaging.

## 2. Materials and methods

### 2.1 Materials

Poly(lactide-co-glycolide) acid (L/G 90/10, M.W. 200,000) was purchased from PolyScitech, Inc (West Lafayette, IN). Dichloromethane (DCM), chloroform, and polyethylene glycol (PEG, Average MW 3350) were purchased from Fisher Chemical (USA). Stearylamine was purchased from Tokyo Chemical Industry CO.,LTD (Tokyo, Japan). Cholesterol was obtained from MP Biomedicals, LLC (Solon, OH). 1,2-distearoyl-sn-glycero-3-phosphoethanolamine-N-[methoxy(polyethylene glycol)-5000 (DSPE-PEG 5000), and 1,2-distearoyl-sn-glycero-3-phosphocholine (DSPC) were purchased from Avanti Polar Lipids, Inc. (Alabaster, AL). Gold-nanorods were purchased from NanoPartz, Inc. (Loveland, CO). Methotrexate (MTX) (Thymoorgan Pharmazie Gmbh, Germany), hydrocortisone and neo-poly-bac ointments were purchased from University of Cincinnati Pharmacy. Sulfo-Cy5 carboxyl acid (Cy5) was purchased from Lumiprobe Co. (Hunt Valley, MD).

### 2.2 Synthesis of MTX/Cy5 encapsulated liposomes

MTX liposomes and MTX/Cy5 co-encapsulated liposomes were synthesized by a modified reverse-phase evaporation (REV) method reported in our previous publication.^8^ MTX liposome was prepared for in vitro drug release test. MTX/Cy5 coencapsulated liposome was used for in vivo drug release test. The MTX or MTX/Cy5 loaded liposome was purified by PD-10 column to remove free drug or dye solution. 50 μL gold nanorod suspension (~4.7×10^13^/mL) (NanoPartz, Loveland, CO) was added in the purified sample, corresponding to a number ratio of 2:1 for a gold nanorod to a liposome. The mixture was gently shaken for at least 40 min to have the gold nanorods attach on the surfaces of liposomes via the electrostatic interaction. The attached gold nanorod-coated liposome suspension was concentrated by centrifugation at 6000 rpm for 30 min to achieve a higher drug concentration (accuSpin Micro 17, Fisher Scientific). After centrifugation, the suspension of concentrated drug/dye loaded liposome (approximate 350 μL) was stored at −20 °C for 2 h and then transferred to −70 °C for another 2 h until completely frozen. Then the frozen liposome suspension was lyophilized in a freeze-dryer (FreeZone 2.5, LABCONCO, Kansas City, MO) for 4 h.

For 1000 μg and 500 μg dosage loading, 36.5 μL and 73 μL of a PBS (phosphate buffer saline) solution, respectively, was added into the lyophilized liposome samples. The rehydrated sample was incubated in room temperature overnight. After stirring the sample with a fine 30-gauge syringe needle was used to manually homogenize the sample for 5 min. Then the gel-like high concentration MTX liposome suspension was centrifuged at 3000 rpm for 3 min to remove any potential bubbles that may occur during the mixing.

The centrifuged samples were diluted 1000 times in PBS. The optical density of diluted samples was measured by UV-vis at 370 nm which was the absorption peak of MTX. The mixed samples were injected into the PLGA implants.

### 2.3 Fabrication of laser-activated drug implant

A laser-activated drug release PLGA implant was generated by loading concentrated MTX liposome or MTX/Cy5 liposome in a PLGA nanoporous tube. The PLGA tube was fabricated by mainly following our previous method.^8^ Briefly, PLGA (MW 200,000, L/G 90/10, PolySciTech) was mixed with polyethylene glycol (PEG) (MW 3350, Fisher Chemical) as porogen at 0.1 PEG to PLGA weight ratio in DCM at 50 mg/ml PLGA in DCM. 800 μl of the mixture was transferred to a mold and bath-sonicated for 80 minutes at 10 °C. The PLGA/PEG solution was air dried overnight in a chemical hood at room temperature and became a ~30 μm thickness sheet. The PLGA/PEG sheet was rolled using a 22-gauge needle as a template twice to create a double-layer tube. One end of the tube was sealed by a preheated iron at 60 °C. 4 μL of the MTX/Cy5 coencapsulated liposome suspension described above was then injected into the one-end sealed tube using a 30-gauge syringe blunt tip needle. The other end of the tube was sealed in the same manner after sample loading. After sealing, the implant was incubated in 1 mL PBS at 35 °C for 24 hours to remove the porogen PEG, resulting in the final product, a nanoporous PLGA capsule implant. Two different liposomal drug doses in the implant, for 1000 μg and 500 μg implants, the same volume 4 uL but dilution was used. For characterization of the dimension of the PLGA/PEG tube, a digital microscope VHX-2000 (Keyence, IL) was used.

### 2.4. In vitro NIR laser irradiation of implant

A laser-activated implant was placed in a 1.5 mL test tube with 1 ml PBS at 35 °C. Each test tube was held horizontally on a petri dish. A pulsed NIR (1064 nm) laser (BrightSolutions, Pavia, Italy) was setup above the test tube. The pulsed NIR laser was irradiated for 15 min total, 5 s at 4 different spots (20 s) with 1 min cooling time between 20 s irradiations. The released MTX (drug) outside of the implant was quantitively analyzed by using UV-vis spectrometry at absorption wavelength 370 nm. A calibration curve was prepared for MTX concentration in PBS corresponding to its optical density (O.D). PBS solution was replaced after every 24 hr. O.D was measured before and after each buffer replacement.

### 2.5. Modelling analysis of drug release

Drug release kinetics was analyzed by fitting cumulative drug release data into following models.

#### 2.5.1. Zero order kinetics

Zero order kinetics is that cumulative drug release is linearly proportional to time, regardless of initial dose in the implant. The equation is

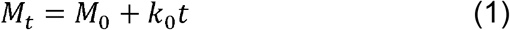

where *M*_0_ is the total drug amount in a surrounding medium after reaching “pleatau”, *M_t_* is the cumulative total amount of drug released in the medium, and *k*_0_ is a zero-order release kinetic constant.

#### 2.5.2. First order kinetics

Drug release rate depends on drug concentration in the drug reservoir, in this case implant. Then,

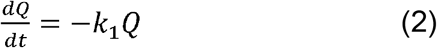

where *Q* is the amount of drug in the implant and *k*_1_ is a first-order release constant. At time *t* = 0, *Q* = *Q*_0_, which is the total amount of drug released from the implant until reaching plateau. Then,

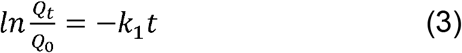

and *Q_t_* = *Q*_0_ – *M_t_*.

#### 2.5.3. Korsmeyer-Peppas (KP) model

The generic equation for the Korsmeyer-Peppas model is

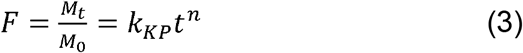

where *F* is fraction of drug released at time *t, M_t_* is drug release from implant at time, *M*_0_ is the total drug release from implant until reaching “plateau”. *k_KP_* is a Korsmeyer-Peppas kinetic constant, and *n* is diffusion or release exponent, which will be determined by fitting. *n* suggests type of drug release mechanism as shown in Table 1.

**Table 1.**
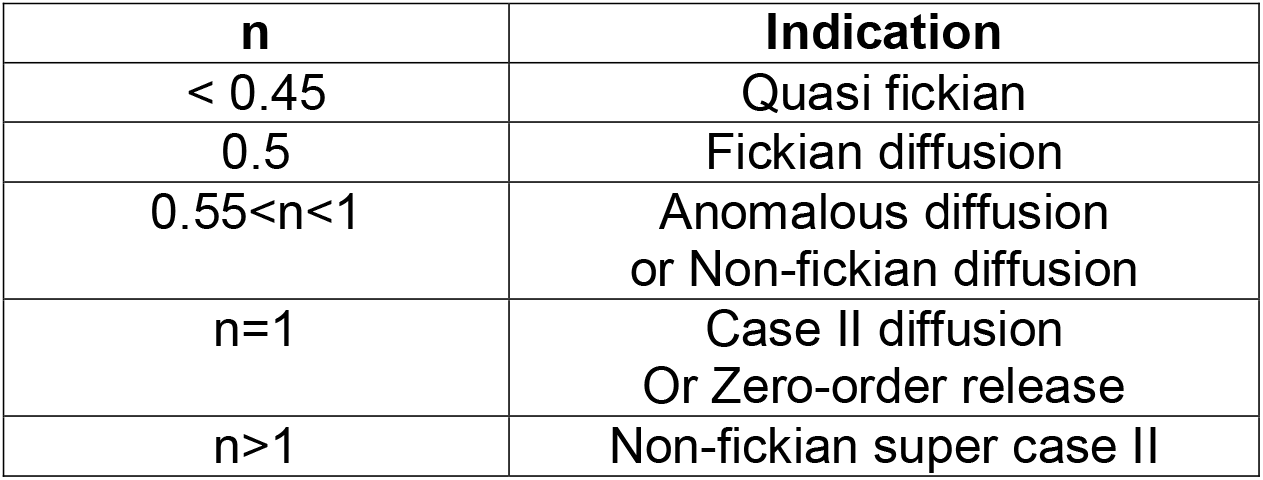
Indication of n value for Korsmeyer-Peppas equation.^10,11^

| **n** | **Indication** |
| --- | --- |
| < 0.45 | Quasi fickian |
| 0.5 | Fickian diffusion |
| 0.55<n<1 | Anomalous diffusion<br>or Non-fickian diffusion |
| n=1 | Case II diffusion<br>Or Zero-order release |
| n>1 | Non-fickian super case II |

The Korsmeyer-Peppas model is applicable to assess the mechanism of the first 60% of drug release taking place from swellable polymeric devices.^11^

#### 2.5.4. Higuchi model

Drug release mechanism primarily caused by diffusion process can be anticipated by fitting the drug release data to the Higuchi release equation as the following.^12^

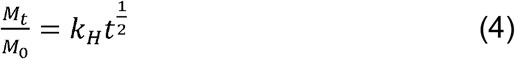

where *M_t_* is drug release from implant at time, *M*_0_ is the total drug release from implant by laser irradiation and *k_H_* is Higuchi constant. It describes drug release as a diffusion process based on Fick’s law.

#### 2.5.5. Statistical analysis of drug release kinetics

Statistical comparisons between release kinetics (*k*_0_, *k*_1_, *k_KP_*, *k_H_*) of 1000 μg implant and 500 μg implant under zero order, first order, KP equation and Higuchi model (with and without first 24 hours data) were performed by Single Factor ANOVA, and p-value <0.05 was considered statistically significantly different (n=3).

### 2.6. Fluorescence calibration of remaining MTX in implants

Implants loaded with 700, 800, 900, and 1000 μg MTX dosage of MTX/Cy5 co-encapsulated liposome suspension were characterized by an in vivo fluorescent microscope (Micron IV, Phoenix Technology Group LLC., CA). The fluorescent intensity of Cy5 dye in the implant was quantified by a mean gray value in ImageJ (NIH). A calibration curve was generated based on the mean gray values in the implant corresponding to the dosage loaded.

### 2.7. In vivo drug release from implant

All animal studies were performed at the Laboratory Animal Medical Services and approved by the IACUC at the University of Cincinnati. The drug release of our laser-activated implant was determined in New Zealand White rabbits (~2 kg). Before injection, the implant was sterilized by ultraviolet (UV) for 30 minutes and inserted into a sterilized 17-gauge needle. The implant was injected into the vitreous at the pars plana through the sclera around 3–5 mm away from the limbus performed by a surgeon. Untreated eyes served as the control group.

The implant in the rabbit eye was examined by optical/fluorescence imaging (Micron IV), ultrasound imaging (Terason Ultrasound, Burlington, MA) and fundus imaging (28D fundus lens, Volk Optical, Mentor, OH). A guide beam was used to aim the center of injected implant. After alignment, a near-infrared pulsed laser (1064 nm, 1.1 W/cm^2^) was replaced with the guide beam. Then the implant was irradiated for 20 s (5 s x 4 different spots along the implant) and halted for 3 min for cool-down. After cooling, irradiation was performed again in the same manner until 10 min total. After the laser irradiation, optical imaging, ultrasound imaging and fluorescence imaging were used to characterize the implant and surrounding area in the vitreous. The lens of microscope was contacted on the cornea of the rabbit with an eye gel (Genteal Tears, Fort Worth, TX). The amount of drug released in vivo was quantified by analyzing the fluorescent intensity of the implant using a mean gray value in ImageJ. Based on the calibration curve obtained in Section 2.6, drug concentration in the implant remained after irradiation was determined. The control eyes did not receive laser irradiation and were imaged in the same manner.

### 2.8. Tissue processing and histological examination

Dissection of the eyes was immediately performed in a sterile biosafety fume hood after eye enucleation. A short incision was carefully made near the limbus using a sharp razor blade to separate the anterior and posterior segments of the eye. Gross images of the posterior segment were taken to show the original position and appearance of the implant. About a quadrant of the posterior segment which was closest to the implant was carefully separated by fine scissors and a razor blade for histological examination. The control eye remained unopened. The isolated segment for the implant+laser group and the control eye were fixed in Davidson’s fixative solution (comprised of 3 parts of 95% ethanol, 2 parts of 37% formaldehyde, 3 parts of distilled water and 1 part of glacial acetic acid) for 24 hours at room temperature. Then they were transferred to freshly prepared 4% paraformaldehyde solution at room temperature until dehydration. The tissue was dehydrated in graded ethanol (50%–100%), cleared in CitriSolv, and subsequently embedded in paraffin. Sequential sections with 8 μm thickness were cut, collected and stained with hematoxylin and eosin (H&E). The histological images were taken by a Nikon Eclipse E800 microscope (Nikon, Tokyo, Japan).

## 3. Results

### 3.1. Characterization of laser-activated drug implant

The PLGA tube prepared from the PLGA/PEG sheet showed the thickness (d), 59.0 ± 7.2 μm, and the inner radius (R), 293.2 ± 2.3 μm (Figure 2, middle). The implant where the lumen was filled with 4 μL of rehydrated MTX/Cy5 liposomes was inserted in a syringe needle for intravitreal injection (Figure 2, right). The concentrations of liposomal MTX in the implant for 1000 μg and 500 μg implants were 245 ± 9.9 mg/ml and 119.5 ± 13.4 mg/ml, respectively, determined by UV-Vis spectroscopy.

**Figure 2.**
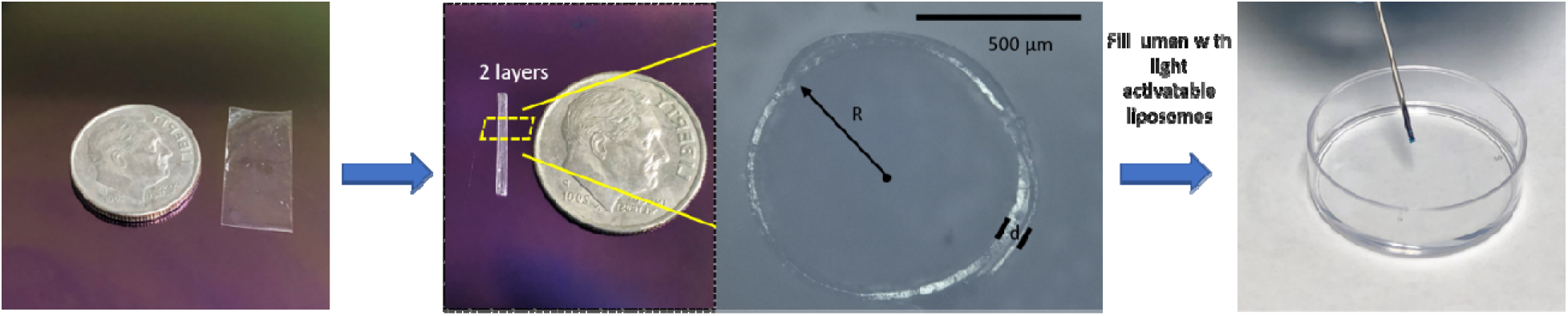
Process of preparation of dye/drug co-encapsulated liposome loaded in a two-layers PLGA implant.

### 3.2 In vitro drug release of implant after laser irradiations

Cumulative amounts of MTX released from the 1000 μg and 500 μg implants after the laser irradiation trigger are demonstrated in Figure 3. Implant initially loaded with 1000 μg implant showed a rapid release within first 24 hr. The release kinetics slope decreased after 24 hr and it reached almost a plateau after 168 hr. For 500 μg implant, the profile was relatively linear with time. The plateau showed up at 192 hours. Overall, ~50 μg and ~25 μg of MTX were released from the 1000 μg and 500 μg, respectively, per irradiation. The results suggest that the “total” dose released by laser was proportional to the initial dose packed in the implant, in other words, the drug concentration.

**Figure 3.**
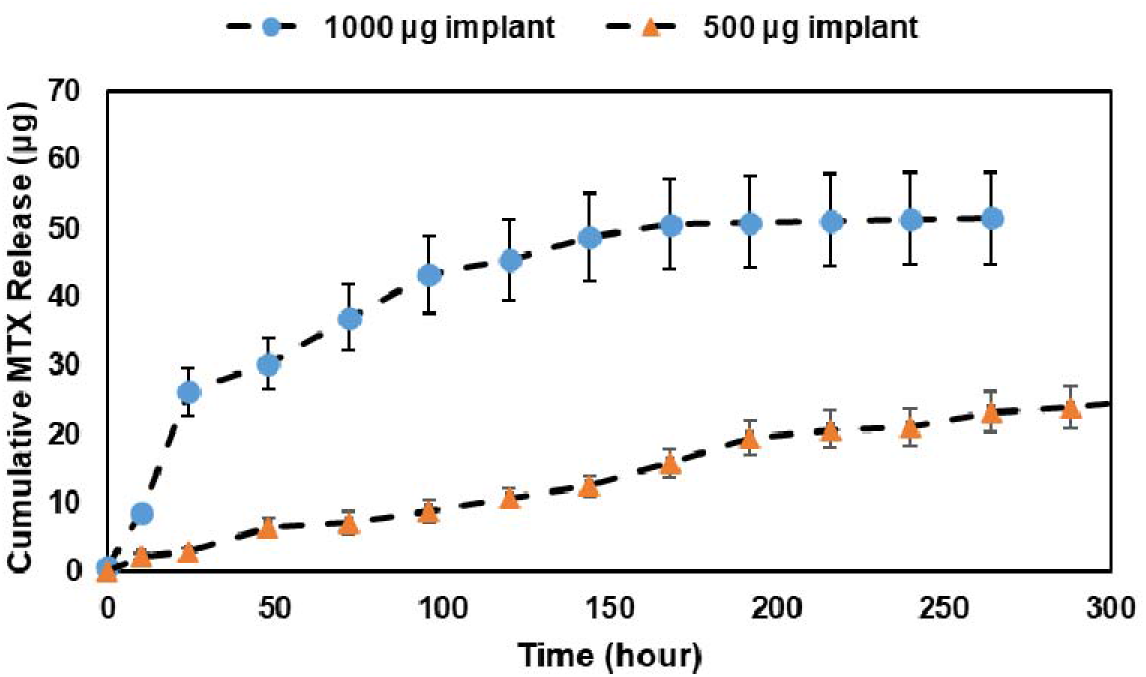
Cumulative amount of MTX released from implants after irradiation with two different total doses as in liposomal form packed in the implant: 1000 μg (blue circles) and 500 μg (orange triangles).

### 3.3 Drug release kinetics models

Cumulative MTX release data were analyzed by fitting into four drug release kinetics models, including zero-order, first-order, Korsmeyer-Peppas, and Higuchi models (Figure 4). For the zero-order fittings, the 500 μg implants showed a higher regression coefficient (R^2^=0.99) than the 1000 μg (R^2^=0.90). (Figure 4(A), Table 2). In fact, the 500 μg kinetic curve fitted the zero-order kinetics the best among the four models. The zero-order rate constants were *k*_0_ =0.0044 and *k*_0_ =0.0036 for 1000 μg and 500 μg, respectively, indicating that the 1000 μg implants released the drug through the capsule membrane faster than the 500 μg. Because of the fast increase of drug release within 24 hr for the 1000 μg implants, we have fitted the data excluding data points before 24 hr. Regardless, the R^2^ values did not improve in the zero-order kinetics for the 1000 μg (R^2^ = 0.82). For the first-order fitting, the 1000 μg drug release kinetic curve fitted the best among the four models (R^2^=0.99), suggesting that the rate of drug release from 1000 μg implants was proportional to the drug concentration released by laser in the implant (Figure 4(B), Table 2). The R^2^ for the 500 μg implants in the first-order kinetic model was 0.95. The first-order kinetic rate constants for 1000 μg and 500 μg were *k*_1_ =0.022 and *k*_1_ =0.0089, respectively, indicating that the drug release rate of the 1000 μg was much faster than the 500 μg.

**Figure 4.**
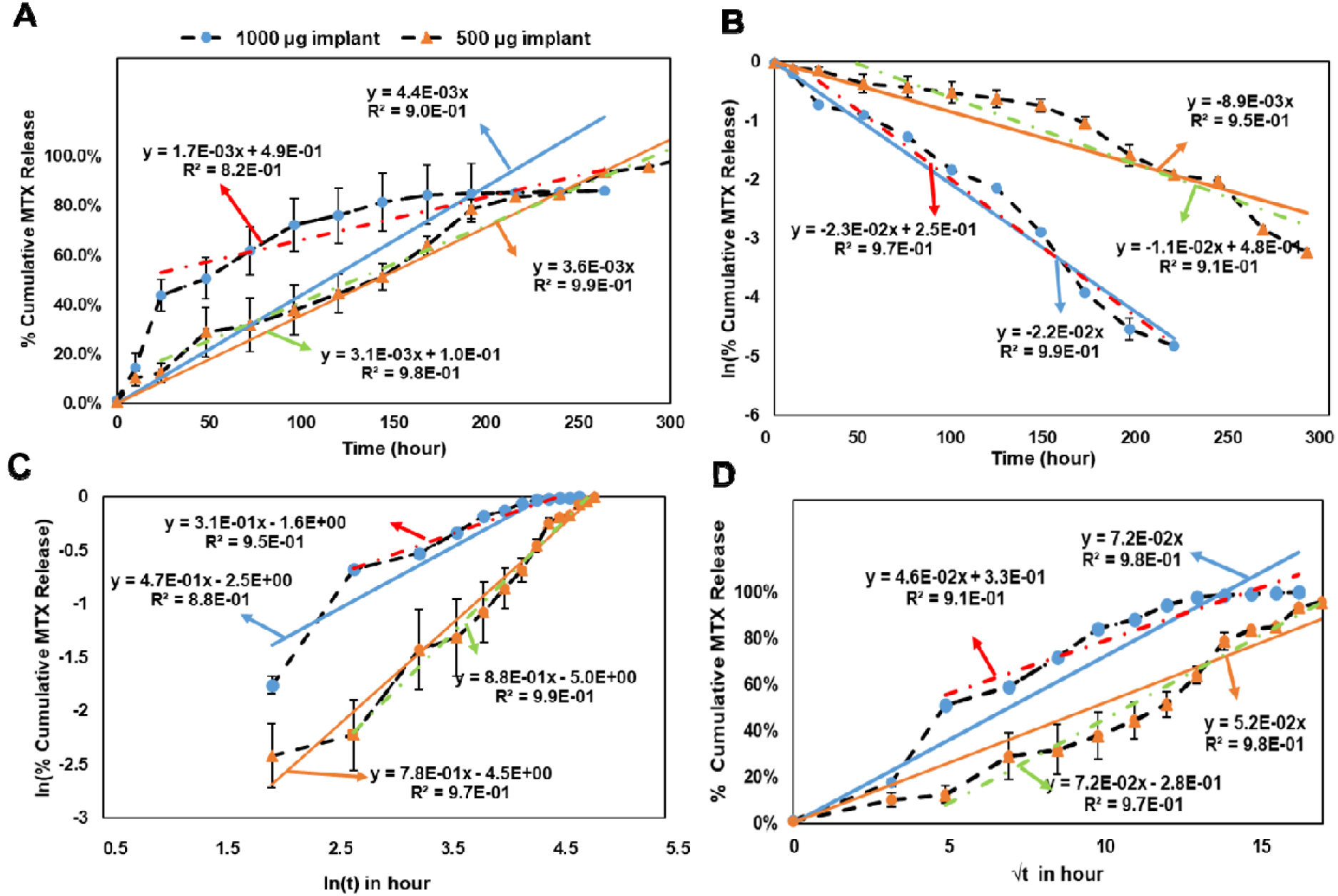
Fitting results of cumulative MTX release kinetics by laser irradiation according to (A) zero-order (B) first-order (C) Korsmeyer-Peppas and (D) Higuchi models. Data: 1000 μg (blue circles) and 500 μg (orange triangles). Trend lines: 1000 μg from 0 hr (blue solid), 1000 μg from 24 hr (red dash-dot), 500 μg from 0 hr (orange solid), and 500 μg from 24 hr (green dash-dot).

**Table 2.** p-values of drug release kinetics for 1000 μg and 500 μg implants.

| Fitting models | $k$<br>(from 0 hr) | $R^2$<br>(from 0 hr) | $k$<br>(from 24 hr) | $R^2$<br>(from 24 hr) | p-values<br>(from 0 hr) | p-values<br>(from 24 hr) |
| --- | --- | --- | --- | --- | --- | --- |
| Zero order<br>(1000 µg) | $k_0 = 0.0044$ | 0.90 | $k_0 = 0.0017$ | 0.82 | 0.32 | 0.048 |
| Zero order<br>(500 µg) | $k_0 = 0.0036$ | 0.99 | $k_0 = 0.0031$ | 0.98 | | |
| First order<br>(1000 µg) | $k_1 = -0.022$ | 0.99 | $k_1 = -0.023$ | 0.97 | 0.00060 | 0.0016 |
| First order<br>(500 µg) | $k_1 = -0.0089$ | 0.95 | $k_1 = -0.011$ | 0.91 | | |
| KP model<br>(1000 µg) | $n_{KP} = 0.47$<br>$k_{KP} = 0.085$ | 0.88 | $n_{KP} = 0.31$<br>$k_{KP} = 0.2019$ | 0.95 | 0.091 ($n_{KP}$)<br>0.024 ($k_{KP}$) | 0.045 ($n_{KP}$)<br>0.0024 ($k_{KP}$) |
| KP model<br>(500 µg) | $n_{KP} = 0.78$<br>$k_{KP} = 0.011$ | 0.97 | $n_{KP} = 0.88$<br>$k_{KP} = 0.0067$ | 0.99 | | |
| Higuchi model<br>(1000 µg) | $k_H = 0.072$ | 0.98 | 0.046 | 0.91 | 0.0044 | 0.066 |
| Higuchi model<br>(500 µg) | $k_H = 0.052$ | 0.98 | 0.072 | 0.97 | | |

For KP model in Figure 4(C), since log(t) at t=0 cannot be plotted, the first data point started from 12 hr. The n values from KP model fittings for 1000 μg and 500 μg were 0.47 and 0.78, respectively, suggesting that they showed different diffusion mechanisms according to Table 1. n=0.47 for the 1000 μg suggested that the drug release from 1000 μg followed Fickian diffusion although the R^2^ value was low (R^2^=0.88). n=0.78 for the 500 μg suggested it followed non-Fickian diffusion with relatively high R^2^ values, at R^2^ =0.97 and R^2^ = 0.99, including data points before 24 hr and excluding 24 hr, respectively. However, the Higuchi model fitting in Figure 4(D) depicts Fickian diffusion, and R^2^ values for both 1000 μg and 500 μg were high at 0.98 and 0.98, respectively, indicative of Fickian diffusion-controlled release. The slopes from Higuchi model showed that *k_H_* =0.072 and *k_H_* =0.052 for 1000 μg and 500 μg, respectively, suggesting drug release from the 1000 μg was faster than the 500 μg.

The KP model originally describes different diffusion mechanisms stem from changes in polymer state or morphology upon hydration.^10^ However, our implant was incubated and equilibrated in an aqueous solvent (PBS) before irradiation. It is unlikely polymer morphology changes from glass to rubber phase due to “swelling” during the drug release. Instead, as the KP model suggests diffusion in glassy polymers may be either Fickian or non-Fickian, depending on the relative rates of chain relaxation and diffusion, and on the activity of water in the polymer. PLGA is glassy compared to the polymers that Korsmeyer and Peppas used, such as PVA hydrogel. Overall, both KP model and Higuchi model suggest drug release mechanism follows diffusion through solid structure.

To confirm the differences in the release of MTX from the 1000 μg and 500 μg implants tested, statistical analysis by Single Factor ANOVA was performed (Table 2). The p-values of most groups were less than 0.05 except zero-order from 0 hr, KP model from 0 hr, and Higuchi model from 24 hr, indicating the rate constants and n values were significantly different between the 1000 μg and the 500. The p-value for the zero-order rate constant *k*_0_ from 0 hr was 0.32. The *k*_0_ was sensitive to the final cumulative drug release amount. Relatively large variations in the 1000 μg group %cumulative release, from 37% to 62%, may have affected the large p-value. The p-value for the KP model from 0 hr was 0.091 and the big error bars in the initial data points of the 500 μg group indicate that the variation could have caused the large p-values. Overall, the statistical analysis implies that each model emphasizes different release mechanisms.

### 3.4 In vivo drug/dye release after laser irradiation

The in vivo study involved intravitreal injection of the implant into the vitreous cavity of rabbits followed by weekly irradiation of the implant with pulsed laser. In Figure 5, the long cylindrical shape of the implant in the vitreous was observed by ultrasound and optical images for the implant + laser group, as opposed to the control group. The ultrasound images show sagittal views that includes the lens on top of the image and the vitreous and the retina. The implant was positioned diagonally following the direction of injection and the position or location did not change over 21 days. The optical (OP) and ultrasound (UL) images in Figure 5 were taken before laser irradiation. Except the Day 0 image, Days 7, 14, and 21 images showed a smudge of dye near the implant, indicative of dye/drug released from the previous week.

**Figure 5.**
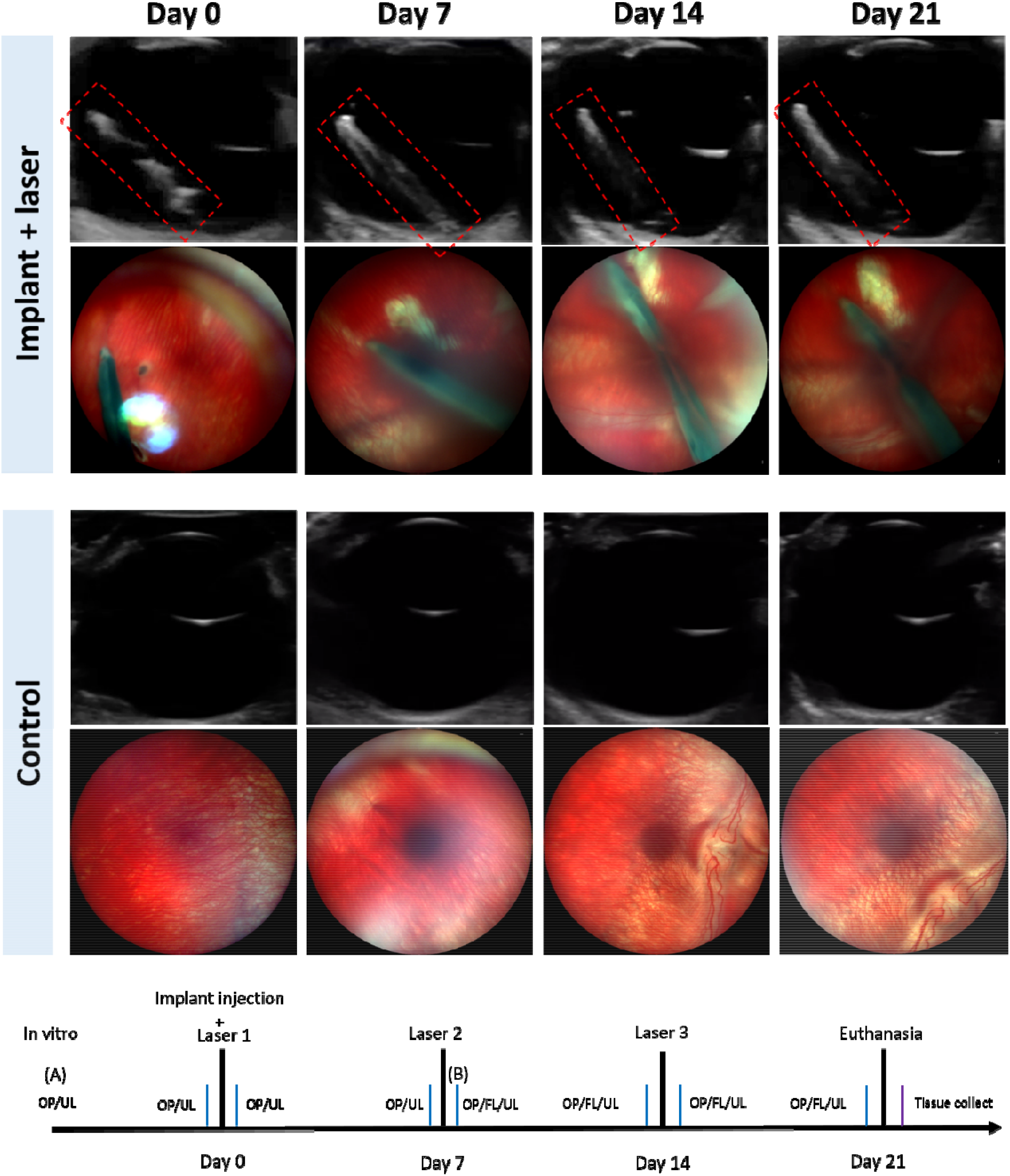
Ultrasound and optical images of the implant-injected eye showing the retina and the implant, and the control eye on Day 0, Day 7, Day 14 and Day 21, respectively. The laser was irradiated at Days 0, 7, and 14 for the implant group.

Based on the in vivo fluorescence images (FL) in Figure 6, the amount of the drug/dye released by laser was estimated. The fluorescence images of the 1000 μg implant before first irradiation and after second irradiations are demonstrated in Figure 6 (A) and (B), respectively. The mean (average) gray values of the whole implant analyzed by ImageJ were 64.1 and 44.6 for (A) and (B), respectively, indicating the dye/drug was released by laser irradiation in vivo. On the other hand, the mean gray values of the whole vitreous increased from 0 to 33.9. Based on the mean gray values in the implant and the calibration curve between the mean gray value in the implant and MTX drug amount in the implant (Supplementary S2), the amounts of MTX in the implant for (A) and (B) were 1000 μg and 898.7 μg, respectively. This suggests that about 100 μg of MTX was released by two irradiations. The results match the in vitro results, where 1000 μg implants released ~50 μg per irradiation. The mean gray values in the center of implant significantly decreased from 84 to 42, indicating that the dye/drug were released from the area where laser irradiated.

**Figure 6.**
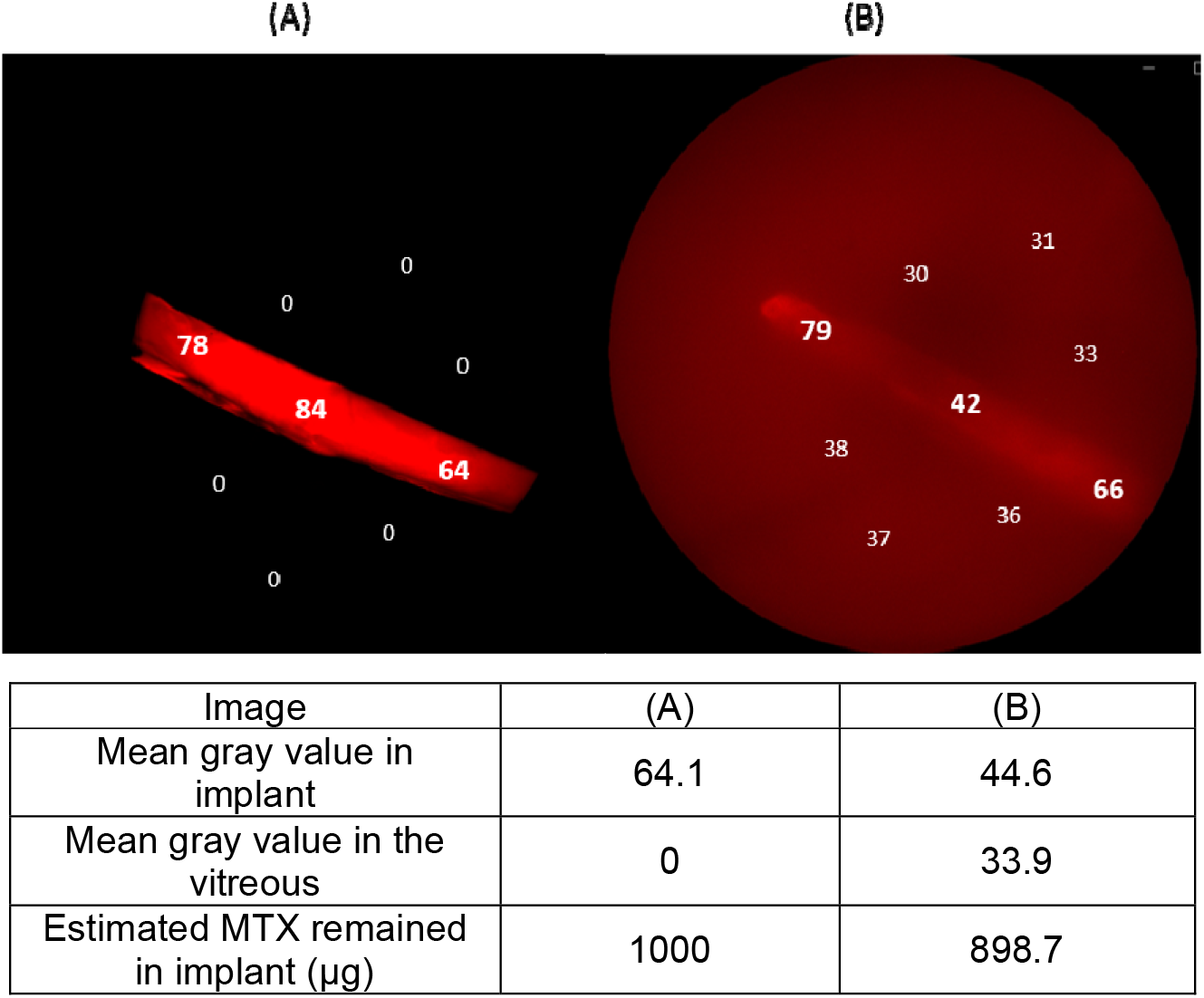
Mean gray values of fluorescent images of (A) before laser and (B) after two lasers in vivo.

### 3.5 In vivo safety of the light-activated drug delivery implant

Based upon the ultrasound and optical images in Figure 5, the implant + laser group did not show any adverse effect on the retina over the 21-day period. The retina margin looked clear and did not change over time in the ultrasound images. Also, in the optical images, abnormal vessels or structures in the retina were not observed. The location and the shape of the implant did not change according to the results of ultrasound and optical imaging.

Figure 7(A) presents the gross picture of the eye cup from the implant+laser group after 21 days and 3 irradiations, showing the intactness of the implant. No obvious damage was observed on the outer surface of the implant. The histological sections from the implant+laser group showed the retinal layers remained their integrity, including nerve fiber layer, inner nuclear layer, outer nuclear layer, retinal pigment epithelium (RPE) and choroid, suggesting minimal toxicity from the implant (Figure 7B). In addition, multinucleated giant cells, which are a typical indicator of foreign body response^13^, were not found in the histological sections. The detachment observed of the retina in the implant eye was due to tissue preparation process, compared to the control eye which was fixed intact without a cut-open for tissue preparation (Figure 8C). Both the optical imaging and histology data showed no sign of laser damages, which are often characterized by RPE and choroidal circular pigmentation, and loss of nuclei in the outer nuclear layer (ONL).^14 15^

**Figure 7.**
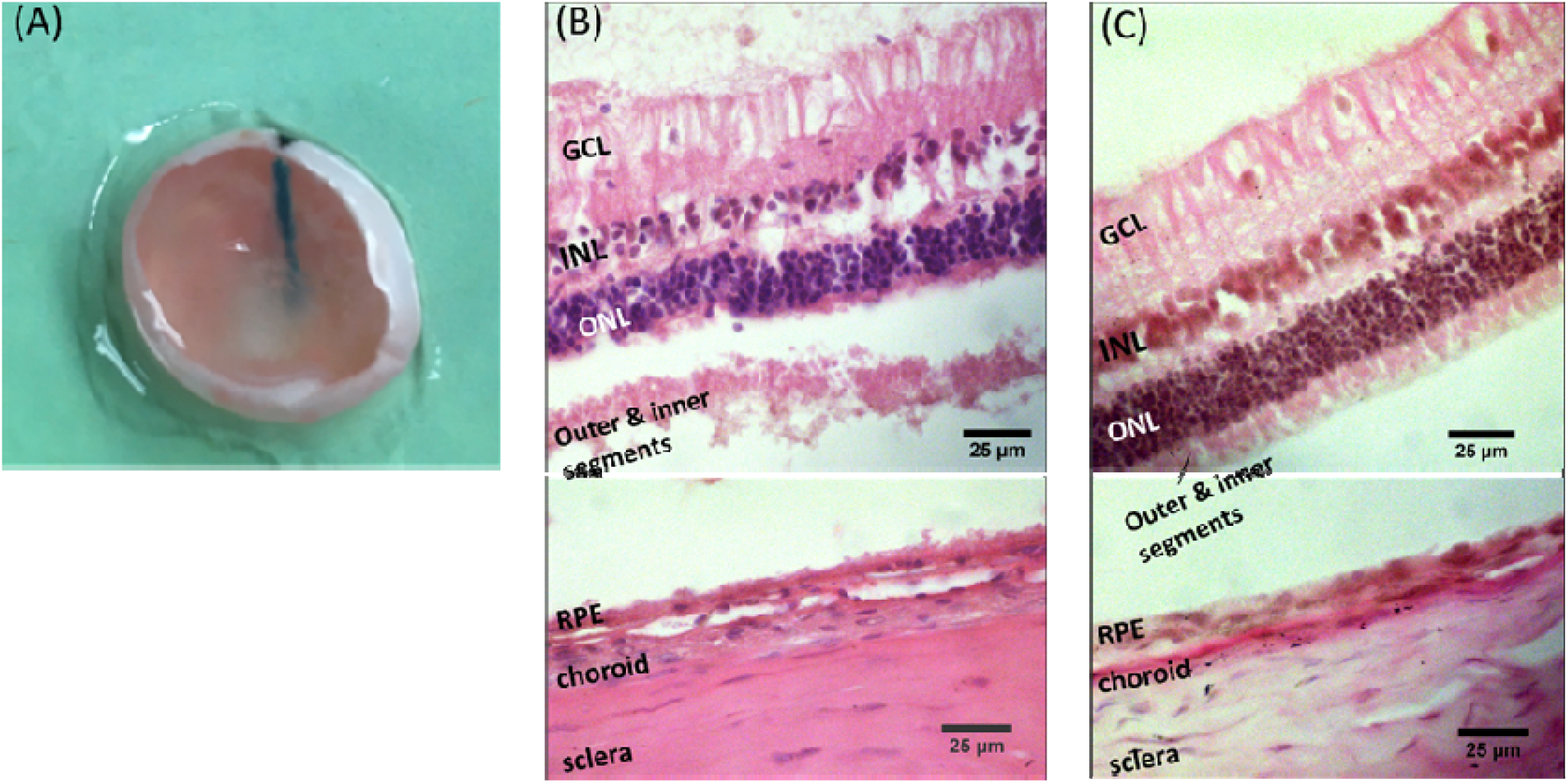
(A) Gross picture of the implant + laser rabbit eye after removing the lens and cornea. Black ink was used to mark the location of implant from outside the sclera. Representative histological images showing the retinal layers with H&E staining of (B) the implant+laser eye and (C) the control eye. GCL: ganglion cell layer; INL: inner nuclear layer; ONL: outer nuclear layer; RPE: retinal pigment epithelium.

## 4. Discussion

Previously, we have tested the long-term stability of the implant for 6 months in vivo and in vitro and the safety in the rabbit eyes.^8^ There were no signs of significant leakage of drug from the implant for 4 months post injection and no cytotoxicity or immune responses were observed after 6 months implantation. In this study, we focused on irradiation induced drug release and achieved a clinically relevant vitreous cavity dose in vivo and in vitro. First, we utilized a lyophilization technique to increase the initial dose of the liposomal drug in the implant compared to our previous study to make the dose relevant to clinical doses. One clinical study showed that 100 ~ 200 μg of MTX per injection biweekly was therapeutically effective to treat proliferative vitreoretinopathy.^16^ Second, the drug dose released per irradiation was also comparable to the clinical dose. Although the total dose of ~50 μg released from the implant after the laser irradiation trigger in this study does not reach the normal clinical dose of 100 ~ 200 μg MTX, the implant provides higher drug concentration in the vitreous over several days than conventional intravitreal injection. This is because the amount of drug in the vitreous after intravitreal injection would significantly drop within a day because of the half-life of free MTX molecules in the vitreous, ~14.3 hr.^17^ In other words, a half of 100 μg, 50 μg, is cleared out from the vitreous in less than a day. On the other hand, the light-activated implant continuously released the 50 μg over 10 days after laser irradiation, keeping a dose of ~5 μg daily. This new delivery mechanism is beneficial for long-term drug delivery. We have also shown repetitive multiple laser-triggered releases in vivo. After 2 irradiations, 100 μg was released from the implant, and theoretically the 1000 μg implant has the capacity for 20 times laser triggers.

The “total” drug release by one laser irradiation was proportional to the total dose packed in the implant: ~50 μg from 1000 μg, and ~25 μg from 500 μg implants. The 1000 μg has double the concentration of liposomal drug compared to the 500 μg implant. Thus, it is likely the higher “total” drug release from the higher dose implants was because of more liposomal drug hit by the laser. In addition, the higher “total” dose released by laser irradiation induced distinct drug release kinetics: the 1000 μg and the 500 μg, respectively, followed first-order kinetics and zero-order kinetics the best. The results imply that the high “total” dose generated a concentration gradient across the implant capsule membrane and the concentration gradient drives the diffusion faster than the low dose. On the other hand, the drug release rate of the low-dose implant was constant, where the concentration gradient did not affect the kinetics. In general, the rate constants of the 1000 μg implants were all higher than the 500 μg, including *k*_0_, *k*_1_, and *k_H_*, indicating the drug release rate of the higher “total” dose was faster than the lower dose.

Both 1000 μg and 500 μg followed Higuchi model with high R^2^ values (R^2^ = 0.98), indicating that drug release followed Fickian diffusion. Higuchi model describes pseudosteady-state diffusion where drug concentration is constant with time throughout the polymer membrane because drug solubility in the membrane is lower than in the reservoir and drug slowly dissolves in the membrane.

Following the mathematical analysis for Fick’s law diffusion for a reservoir cylinder, where drug molecules diffuse from a lumen through a vessel wall,^18^

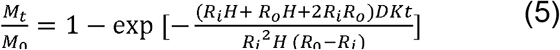

the diffusion coefficients (*D*) of MTX through the membrane were *D* =4.2 x 10^-10^ cm^2^/s and 1.9 x 10^-10^ cm^2^/s, for the 1000 μg and 500 μg, respectively, where *R_i_* is inner diameter, *R_o_* is outer diameter, *H* is the length of the cylinder (implant), and *K* is the partition coefficient of MTX between the membrane and the reservoir. We assume *K* =1 and MTX is penetrating through the pores. We found that the effective diffusion coefficient of MTX through the nanopores was approximately 1000 times less than the diffusion coefficient of MTX in water. In addition, the *D* was proportional to the rate constant of the first-order kinetics *k*_1_ and the drug release rate of the 1000 μg was twice as fast as the 500 μg. These results were consistent with other studies, which described that increasing the loading of therapeutic particles in the matrix also increases the release rate of therapeutics.^19–22^ Also the effective diffusion coefficients based on Fick’s law were in the same order at 10^-10^ cm^2^/s and the values were increasing with the loading capacity.

In vivo drug release was successfully shown via the implant’s fluorescence intensity changes (Figures 5 and 6). The mean gray values in the implant decreased while the value in the vitreous increased after laser irradiation. The laser used in this study was a pulsed picosecond laser with 700 ps pulse duration, 10 kHz repetition frequency at 1064 nm wavelength at 1 W. Although our previous study has shown that drug was released by a single pulse of this laser,^23^ 5 s was used per one spot in order to release physiologically relevant dose in this study. Our other previous study also has shown that the longer laser duration, the more drug is released.^24^ For the safety of the retina, we calculated maximum permissible exposure based on the following equation.^25^

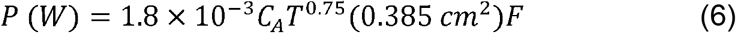

where *C_A_* = 5 for (*λ* = 1064 *nm*), *T* is total irradiation duration, 0.385 *cm*^2^ is area of 7 mm pupil, and *F* is pulse repetition frequency. The calculation suggested that the laser intensity for 5 s at 1 W is 9 times higher than the maximum permissible exposure value with a focal length at 44 mm. Note that this power is calculated when the laser directly hit the retina whereas laser should be targeted to implants. In the future studies, power as low as 100 mW will be used to avoid potential laser damage.

## 5. Conclusion

In this study, we successfully showed effective drug release from a nanoporous PLGA implant using pulsed NIR laser irradiation both in vitro and in vivo. We analyzed the drug release kinetics in vitro by fitting utilizing zero order, first order, and KP and Highchi models. We quantified multiple drug releases in the vitreous in the in vivo fluorescence study, consistent with the in vitro data. The dose released from the implant after the laser irradiation trigger was also clinically relevant. Histology and optical and ultrasound imaging data suggest the drug delivery system is not toxic to the retina. This drug delivery system could be potentially used for long-term posterior eye disease treatment.

## Acknowledgment

This study was partially supported by Ohio Lions Eye Research Foundation, Office of Research at University of Cincinnati, and NIH KL2 award (5KL2TR001426-04). The authors would like to thank Drs. Jonathon Nickels and Vesselin Shanov for equipment.

